# Pre-implantation genetic diagnosis for a family with Usher syndrome through targeted sequencing and haplotype analysis

**DOI:** 10.1101/460378

**Authors:** Haining Luo, Chao Chen, Yun Yang, Yuan Yuan, Wanyang Wang, Renhua Wu, Yinfeng Zhang, Zhiyu Peng, Ying Han, Lu Jiang, Ruqiang Yao, Xiaoying An, Weiwei Zhang, Yanqun Le, Jiale Xiang, Na Yi, Hui Huang, Wei Li, Yunshan Zhang, Jun Sun

## Abstract

**Objective:** Our objective was to investigate the applicability of targeted capture massively parallel sequencing in developing personalized pre-implantation genetic diagnosis (PGD) assay.

**Methods:** One couple at risk of transmitting Usher Syndrome to their offspring was recruited to this study. The genomics DNA (gDNA) was extracted from the peripheral blood and underwent in vitro fertilization (IVF)-PGD. Prenatal molecular diagnosis was performed in the 20th week of gestation and the chromosomal anomaly was analyzed.

**Results:** Customized capture probe targeted at *USH2A* gene and 350kb flanking region were designed for PGD. Eleven blastocysts were biopsied and amplified by using multiple displacement amplification (MDA) and capture sequencing. A HMM-based haplotype analysis was performed to deduce embryo’s genotype by using SNPs identified in each sample. Four embryos were diagnosed as free of father’s rare mutation, two were transferred and one achieved a successful pregnancy. The fetal genotype was confirmed by Sanger sequencing of fetal genomic DNA obtained by amniocentesis. The PGD and prenatal diagnosis results were further confirmed by the molecular diagnosis of the baby’s genomic DNA sample. The auditory test showed that the hearing was normal.

**Conclusion:** Targeted capture massively parallel sequencing (MPS) is an effective and convenient strategy to develop customized PGD assay.

**Key points:** Genetic counseling session was conducted with a family having Usher patient who was molecularly diagnosed, and a healthy baby was born with the help of successful PGD assay. This is of vast importance in management plans since it is the first report of PGD in Usher syndrome based on targeted capture MPS.

## 1 Introduction

Usher syndrome is an autosomal recessive disorder that results in hearing loss and progressive retinitis pigmentosa (RP). It is estimated to account for 50% of combined deafness and blindness in adults [1]. The identification of causative mutations in high-risk families is helpful for genetic counseling and reproductive management. Once pathogenic variants are identified, pre-implantation genetic diagnosis (PGD) can be applied to prevent the conceiving-affected babys. Conventionally, PGD is carried out by directly analyzing the mutation of interest or through short tandem repeat (STR) linkage analysis [2-4]. However, it will be a time-consuming and labor-intensive process to choose informative STRs for the family of interest and to optimize the experiment for mutation analysis.

Benefited from the development of massively parallel sequencing (MPS), the emerging MPS-based approach has been developed and applied for the molecular diagnosis of the single gene disease[5] or chromosomal anomaly[6, 7] in different reproduction stages ranging from preconception to neonate care. The development of MPS also brings a new strategy to develop PGD for single gene diseases. Targeted capture MPS can analyze plenty of SNPs flanking the gene of interest at one time, which can ensure to obtain enough informative SNPs for linkage analysis and to minimize the influence of ADO on the accuracy of PGD. As the process of capture probe design is matured, it would be easy to develop customized PGD assay according to each case’s requirements. The feasibility of targeted capture MPS and linkage analysis based PGD has been proved in a thalassemia family study [8].

In this study, we report the development and successful application of targeted capture MPS and haplotype analysis-based in-vitro fertilization (IVF)-PGD protocol in an Usher syndrome family, coupling with prenatal testing for fetal aneuploidy and large chromosomal imbalance arrangement using an MPS-based method. This is the first report of PGD in Usher syndrome based on targeted capture MPS.

## 2 Materials and Methods

### 2.1 Patient

The patient was a 27-years-old woman who was reported to have the onset of visual loss and night vision loss 11 years ago. Ophthalmologic examinations showed that she suffered from bilateral retinitis pigmentosa (RP), and audiometry tests showed that she had a bilateral sensorineural hearing loss. Her father also suffered from RP and sensorineural hearing loss. Her mother and husband were apparently healthy. The woman was planning pregnancy but was worried about that the child might be affected by the same disease. The patient had a strong desire to find out the cause of the disease and to have an unaffected child via PGD. Approval for this study was obtained from the BGI Board of the Ethics Committee. The written consent was obtained from the participants.

### 2.2 Genetic molecular diagnosis by targeted capture MPS

After carefully genetic counseling, a multiple gene panel assay containing 67 genes associated with Usher syndrome and RP (Supplemental Table 1) was recommended to the family to identify the disease-causing gene. Molecular genetic analysis was performed on the patient and her parents. Her husband’s sample was also analyzed in parallel as a screening test to check whether he carries any mutation in the disease-causing gene.

5 ml EDTA-anticoagulant peripheral blood samples were collected from the patient, her parents, her husband and her parents-in-law, respectively. The genomics DNA (gDNA) was extracted from the peripheral blood using QIAamp DNA Blood Midi Kit (Qiagen, Hilden, Germany). gDNA of the patient, her parents and her husband were captured sequencing using custom-designed NimbleGen SeqCap EZ probes targeted at exons and 10bp flanking sequences of the 67 genes. Libraries were prepared by following the manufacturer’s instruction and sequenced for 101-bp paired-end reads on Hiseq 2500. Then the paired-end sequencing reads were mapped to the human reference genome (Hg19, GRCh37) using BWA MEM (0.7.12) with default parameters. SNV and small indel were called by using GATK. Variants detected in target genes were annotated and interpreted. After the identification of the disease-causing mutations, Sanger sequencing was performed in all family members to confirm the reality and carrier status by using the primers and reaction conditions shown in Supplemental Table 2.

### 2.3 In-vitro fertilization (IVF)

The couple went to Tianjin Central Hospital of Gynecology Obstetrics to undergo an IVF-PGD cycle. The patient underwent an IVF protocol including pituitary down regulation with a gonadotropin hormone releasing agonist. This was then followed by the continued use of the gonadotropin hormone releasing agonist with the addition of a combination of 150 IU of recombinant follicle stimulating hormone (FSH Gonal-F, Merck Serono) and 225 U of human menopausal gonadotropin (HMG, Lizhu China) for a total of 12 days of stimulation. Transvaginal ultrasound monitoring and estradiol measurements were performed to assess follicular maturity with a peak estradiol level of 9556 pg/mL on the last day of stimulation. Human chorionic gonadotropin was administered when there were two lead follicles of 18 mm in average diameter and transvaginal ultrasound guided oocyte retrieval was performed 34 h later. Twenty-four oocytes were recovered and 21 mature oocytes underwent intra-cytoplasmic sperm injection (ICSI). All 21 oocytes fertilized and 11 embryos were subsequently biopsied at the blastocyst stage.

### 2.4 Preimplantation genetic diagnosis (PGD) by targeted MPS

Blastocysts biopsy samples underwent multiple displacement amplification (MDA) using REPLI-g Midi kit (Qiagen) according to the instruction. The MDA products were quantified with Qubit fluorometric (Thermo Fisher). MDA products of the 11 embryos and gDNA of the patient, her parents, her husband and her parents-in-laws were all capture based sequenced with a customized hybridization probe (CustomArray) designed to enrich 1.5 Mb region covering 350kb upstream to 350kb downstream of *USH2A* gene. The sequencing libraries were prepared with 1 μg MDA products or gDNA according to the manufacturer’s instruction. Then the libraries were pair-end sequenced by Hiseq 2500 with a read length of 101 bp.

After low-quality reads were removed, the remaining reads were aligned to the human reference genome (hg19, build 37) by using BWA software (0.7.12) with default parameters. The removal of PCR duplication reads and verification of mate-pair information were performed with Picard (1.87). Then, the SNP calling was performed with the GATK software. The SNPs identified in the couple and their parents were used for haplotype construction. The couple’s haplotypes and their associations with mutations of interest were constructed through a parents-child trios’ strategy according to Mendel’s law.

Allele drop out (ADO) is randomly happened across the genome during MDA, which can result in amplification failure of one of two heterozygous alleles in the embryo and may result in the misdiagnosis of embryo’s genotype. We evaluated ADO rate for each embryo sample by using alleles that were different homozygous in the couple (e.g., mother AA, father BB), and the ADO rate can be calculated as the percentage of allele identified as homozygous in the sample.

In order to avoid the influence of ADO in the embryo’s genotype analysis, only heterozygous alleles identified in the embryo sample were used for linkage analysis. Maternal haplotype inheritance was deduced with an allele that is homozygous in father and heterozygous in mother. Paternal haplotype inheritance was deduced with an allele that is homozygous in mother and heterozygous in father. A hidden Markov model (HMM) was built based on the allele status in each SNP, and the fetal inheritance haplotypes were deciphered with the Viterbi algorithm as Xu et al. previously described[8].

### 2.5 Prenatal diagnosis

Prenatal molecular diagnosis was performed in the 20^th^ week of gestation by using genomic DNA extracted from 20 mL amniotic fluid. Fetal genotype was confirmed by PCR and Sanger sequencing of amniotic fluid DNA. Aneuploidy and chromosomal imbalanced arrangements larger than 1 Mb were detected with low-coverage whole-genome NGS approach. Briefly, 100 ng genomic DNA extracted from the amniotic fluid sample was sheared into small fragments ranging from 100 to 400 bp with Bioruptor (Diagenode). After end-repair, “A” overhanging and adapter-ligation, DNA fragments underwent 12 cycles of PCR with multiplex primers. PCR products were purified with Agencourt AMPure Kit (Beckman). The library was sequenced with single-end 50-cycle sequencing on Hiseq2000 with the data production of 0.75 Gb. The chromosomal anomaly was analyzed as Dong et al described[5, 6, 7].

### 2.6 Genetic and auditory examinations of newborns

2ml EDTA-anticoagulant peripheral blood was collected from the newborn for Sanger sequencing to further confirm the PGD and prenatal diagnosis result. And the auditory examination was performed 72h later after birth by ear acoustic emission analyzer (Maico eroscan).

## 3 Results

### 3.1 The results of genetic testing by target capture sequencing

In order to identify causative variants in the family, the capture sequencing of exons of genes associated with Usher syndrome and RP were performed for the patient, her parents and her husband. mean depth of 130-fold per family member was obtained, with an average 91.3% of the region covered with at least 20 reads (Supplemental Table 3). Compound heterozygous c.1144-2A>C and c.6752C>A mutations in the *USH2A* gene were detected in the patient. Both mutations were pathogenic mutations with the allele frequency of zero in dbSNP, HapMap and 1000 Genome database. c.1144-2A>C is in the splice acceptor site which probably affects the splicing process. c.6752C>A (p. Ser2251Ter) is a nonsense mutation which introduces a terminational codon in 2251 amino acids. The family analysis revealed that the c.1144-2A>C mutation was inherited from the patient’s mother who was a carrier of this heterozygous mutation, while the mutation c.6752C>A (p.Ser2251Ter) was inherited from the patient’s father who was compound heterozygous for the nonsense mutation c.6752C>A (p. Ser2251Ter) and a known pathogenic missense mutation c.9815C>T (p.Pro3272Leu) [9, 10]. Thus, the causative mutation was successfully identified in the family. Both the patient and her father were diagnosed as Usher Syndrome IIA.

The targeted sequencing of the patient’s husband showed that he was heterozygous for c.10740+7G>A mutation in *USH2A*. c.10740+7G>A is a rare mutation with an allele frequency of 0.001 in dbSNP. It is a variant of unknown significance, but the result predicted by MutationTaster showed that it could form a new donor site. After carefully genetic counseling, the couple hoped to take an IVF and PGD cycle to make sure that their children do not inherit the c.10740+7G>A mutation to minimize the risk of being affected by Usher syndrome. The carrier status of each family member was further validated with Sanger sequencing. (Fig. 1 and Supplemental Fig. 1)

**Fig. 1.**
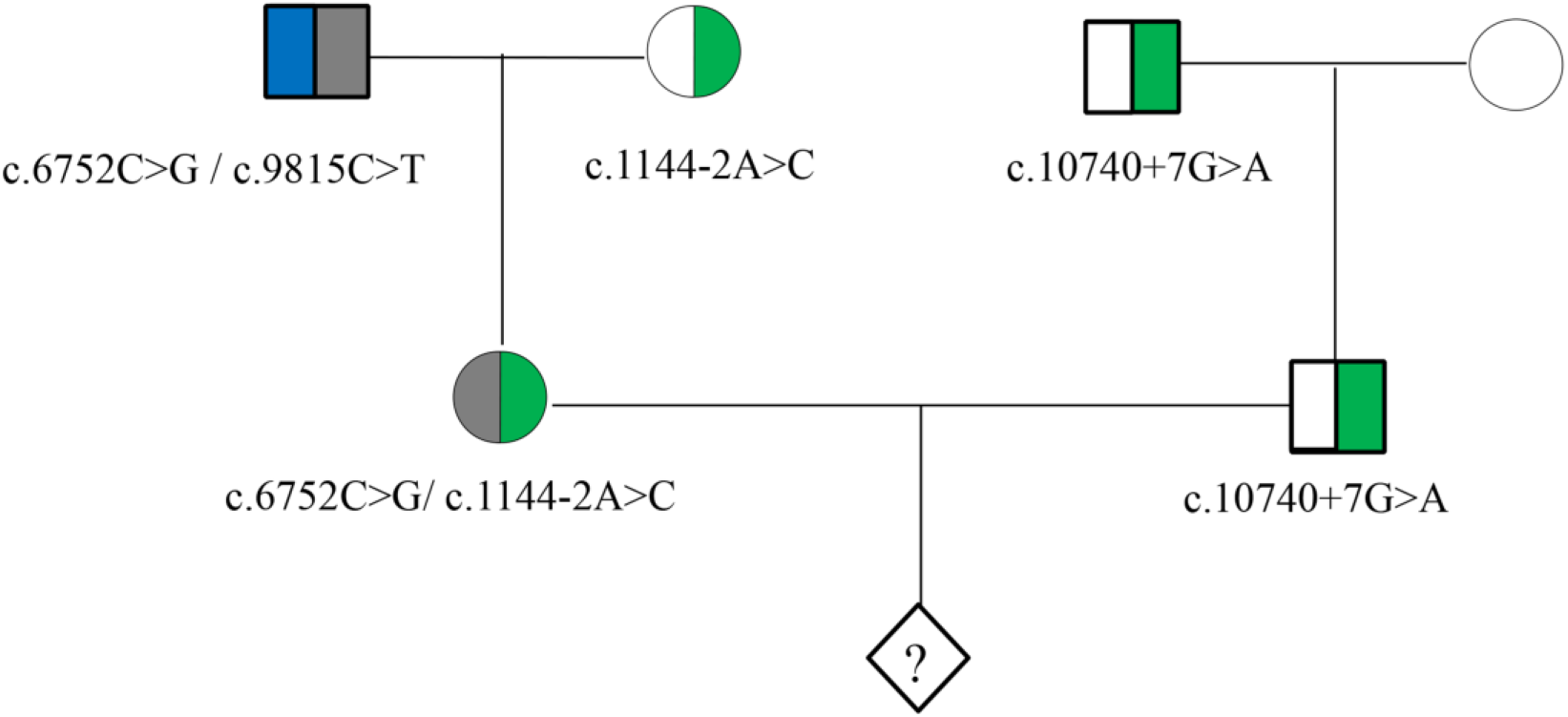
Pedigree of the family. Half-shaded areas indicate carrier state, rhombus indicates embryo.

### 3.2 Haplotype analysis of *USH2A* gene and 350kb flanking region in the couple

To construct the couple’s haplotype, gDNA obtained from the patient, her parents, her husband, and her parents-in-law were capture sequenced for the region covering 350kb upstream *USH2A* to 350kb downstream *USH2A* gene. A mean depth of 411-fold per family member was obtained (Supplemental Table 4). The haplotypes were established with SNP information on the targeted region of the couple and their parents according to Mendelian laws. We defined haplotype linked to c.6752C>A as M-Hap A, while haplotype linked to c.1144-2A>C as M-Hap B in the patient. For the husband’s haplotype, we defined haplotype linked to c.10740+7G>A as F-HapA, and haplotype linked to wildtype allele as F-HapB. We defined SNP as informative if it was only present in one haplotype. An average of 2883 SNPs were identified in each family member.1115 and 1320 SNPs were successfully phased in the patient and her husband, respectively.184 SNPs were identified as heterozygous in the patient but homozygous in her husband and could be used for maternal haplotype inheritance deduction, in which 101 SNPs were M-HapA informative and 83 SNPs were M-HapB informative. 483 SNPs were identified as homozygous in the patient but heterozygous in her husband and could be used for paternal haplotype inheritance deduction, in which 191 SNPs were P-HapA informative and 292 SNPs were P-HapB informative.

### 3.3 Genetic diagnosis in clinical PGD cycle

A mean depth of 189-fold (range: 136-313) was obtained for blood samples and embryo biopsies respectively, with an average 82.05% of the region (range: 68.68%-88.24%) covered by at least 30 reads (Supplemental Table 4). An average of 41 different homozygous SNPs covered by at least 30 reads in the couple were used for ADO rate calculation. The ADO rate was ranged from 0% to 31.7% in different embryos.

Haplotype inheritance deduction in embryo biopsy sample was carried out by using only heterozygous SNPs identified in each embryo to prevent the mistake that might be caused by ADO. An average of 260 informative heterozygous SNPs (range: 111-379) were identified in embryo biopsy samples, of which an average of 186 SNPs (range: 77-282) were used to deduce paternal inherited haplotype and 73 SNPs (range: 34-97) were used to deduce maternal inherited haplotype. The genotypes of the 11 embryos were all successfully deduced by using the HMM approach. Four embryos were free of paternal rare *USH2A* mutation, embryos 1, 8, 11 were carriers of p.Ser2251Ter mutation and embryo 9 was a carrier of c.1144-2A>C. While other embryos were all compound heterozygous, embryos 2, 3, 4, 5, 6, 10 were compound heterozygotes of c.1144-2A>C and c.10740+7G>A, and embryo 7 was a compound heterozygote of p.Ser2251Ter and c.10740+7G>A. (Table1, Fig. 2)

**Fig. 2.**
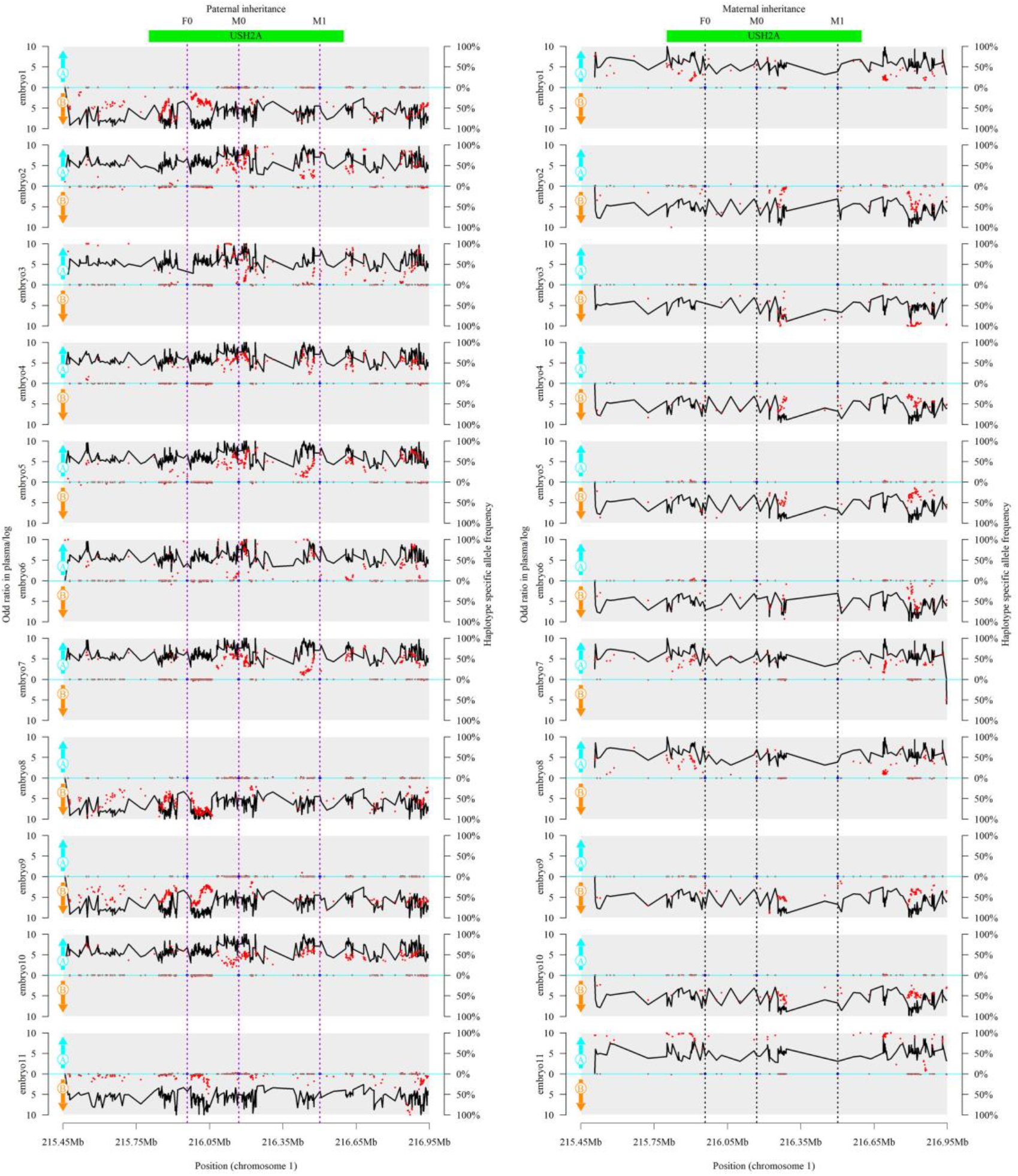
PGD haplotype analysis in embryo 1 to 11. Left, embryo inheritance of paternal haplotype. Right, embryo inheritance of maternal haplotype. The X-axis represents the loci on chromosome 1. The red points represent the allele frequencies of haplotype informative alleles among plasma reads, haplotype A specific allele was drawn above X-axis, and haplotype B specific allele was drawn below X-axis. The black line represents the logarithmic values of the odd ratios of the inherited haplotype. The paternal haplotype A carries c.10740+7G>A mutation (F0), the paternal haplotype B is wild type. The maternal haplotype A carries p.Ser2251Ter (M0), and the maternal haplotype B carries c.1144-2A>C mutation (M1). The vertical dot line indicates the location of target mutation.

**Table 1.** The haplotype in *USH2A* gene in 11 embryos. M-Hap A: p.Ser2251Ter; M-Hap B: c.1144-2 A>C; F-Hap A: c.10740+7G>A; F-Hap B: wild type

| Embryo | ADO | Haplotypes in *USH2A* gene | Genotypes in *USH2A* gene | Numbers of Informative SNPs supported each haplotype in each embryo |  |  |  |
| --- | --- | --- | --- | --- | --- | --- | --- |
|  |  |  |  | M-Hap A | M-Hap B | F-Hap A | F-Hap B |
| Embryo 1 | 0.00% | M-Hap A/F-Hap B | p.Ser2251Ter/N | 97 | 0 | 0 | 282 |
| Embryo 2 | 0.00% | M-Hap B/F-Hap A | c.1144-2A>C/c.10740+7G>A | 0 | 66 | 168 | 0 |
| Embryo 3 | 31.71% | M-Hap B/F-Hap A | c.1144-2A>C/c.10740+7G>A | 0 | 32 | 118 | 0 |
| Embryo 4 | 0.00% | M-Hap B/F-Hap A | c.1144-2A>C/c.10740+7G>A | 0 | 82 | 183 | 0 |
| Embryo 5 | 0.00% | M-Hap B/F-Hap A | c.1144-2A>C/c.10740+7G>A | 0 | 82 | 185 | 0 |
| Embryo 6 | 12.20% | M-Hap B/F-Hap A | c.1144-2A>C/c.10740+7G>A | 0 | 58 | 117 | 0 |
| Embryo 7 | 0.00% | M-Hap A/F-Hap A | p.Ser2251Ter/c.10740+7G>A | 94 | 2 | 185 | 0 |
| Embryo 8 | 4.88% | M-Hap A/ F-Hap B | p.Ser2251Ter/N | 95 | 0 | 0 | 276 |
| Embryo 9 | 0.00% | M-Hap B/F-Hap B | c.1144-2A>C/N | 0 | 83 | 0 | 282 |
| Embryo10 | 0.00% | M-Hap B/F-Hap A | c.1144-2A>C/c.10740+7G>A | 0 | 81 | 181 | 0 |
| Embryo11 | 14.63% | M-Hap A/F-Hap B | p.Ser2251Ter/N | 34 | 0 | 0 | 77 |

### 3.4 Transfer of Embryos and prenatal diagnosis

At the first time, embryo 9 was thawed and transferred on the scheduled day according to PGD results. Frozen-thawed blastocyst transfer was performed in a hormone replacement cycle combining external estrogen pills. The transfer was carried under ultrasound guidance when endometrium thickness reached 7.5 mm. But the embryo failed to implant. Four months later, embryo 1 was transferred with the same protocol when endometrium thickness reached 7.8 mm. A positive β-HCG was obtained 14 days after embryo transfer and fetal heart beat was confirmed by ultrasound observation 4 weeks after embryo transfer.

In the 20th week of gestation, the amniotic fluid was collected for confirming the fetal genotype with Sanger sequencing and testing chromosome imbalance anomaly with NGS. Sanger sequencing result showed that the fetus was a carrier of p.Ser2251Ter. This proves the accuracy of PGD (Fig. 3a). The chromosome imbalance anomaly results showed that no CNV larger than 100kb was identified in the fetus (Fig. 3b).

**Fig. 3.**
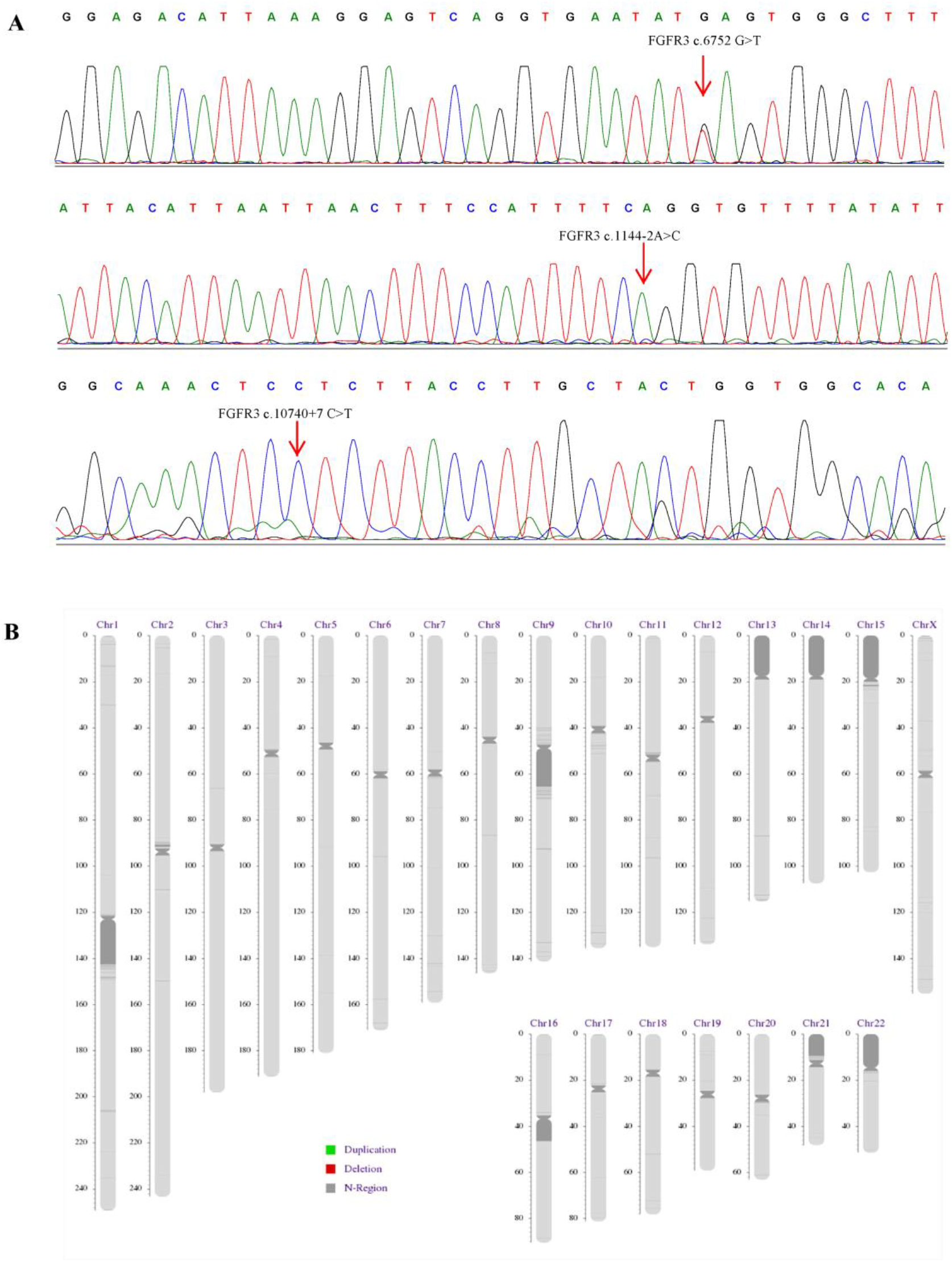
Prenatal diagnosis of amniotic fluid DNA. a. Sanger sequencing results of amniotic fluid DNA confirmed the PGD results, showing the heterozygous c.6752C>A mutation and wild type of c.1144-2A and c.10740+7G. And b. chromosome imbalance anomaly results showed that no CNV larger than 100kb was identified in the fetus.

### 3.5 Genetic and auditory examination after birth

A female baby weighting 2850 g was delivered in the 38th week of gestation, having apparently normal phenotypes. The PGD and prenatal diagnosis result were confirmed with the molecular diagnosis of the baby’s cord blood (Supplemental Fig. 2). The auditory examination result was normal.

## 4 Discussion

Usher syndrome is an autosomal recessive disorder and when both parents are carriers for Usher syndrome, each child has a 1 in 4 (25 percent) chance of inheriting the two changed gene copies, but the risk increase to 1 in 2 if one of the parents is a patient. PGD is an effective method to prevent the conceiving of affected children.

Here, we present an MPS based haplotype linkage PGD analysis for an Usher syndrome-affected family. After the disease-causing mutation was identified, a personality designed PGD targeted at *USH2A* was developed and applied to this family. In addition, MPS-based invasive prenatal detection of fetal aneuploidy and large chromosomal imbalance arrangement were also performed to ensure that the fetus was normal.

The identification and screening of the causative genetic variant in high-risk families is an important precondition for pre-implantation genetic diagnosis. It has been reported that every person carries an average of 2.8 recessive mutations [11]. In the above family, the patient and her father both were compound heterozygous of pathogenic mutations in the *USH2A* gene and were diagnosed as Usher syndrome. The patient inherited one pathogenic mutation from her carrier mother. This also emphasizes the importance of screening the partner of the recessive disease patient to evaluate the risk of conceiving children affected by the same disease. Thus, the same gene panel was applied to the patient’s husband for carrier testing, and a rare mutation (c.10740+7G>A) was detected in the *USH2A* gene. Although the clinical significance of c.10740+7G>A is not clear, this variant was predicted to affect the splicing process by mutation taster, so the pathogenicity can’t be excluded. Meanwhile, after careful counselling, the couple hoped to receive IVF-PGD to prevent the embryo from inheriting c.10740+7G>A mutation from the husband. The identification of the mutation carrier status in each family member provides the genetic information needed for PGD implementation.

The linkage analysis of genetic markers is an important approach to prevent misdiagnosis that may be caused by ADO in PGD. In the past decades, STR is a frequently used genetic marker in PGD, but it may be time-consuming and labor-intensive to select appropriate markers for the family of interest and to optimize the experiments. The capture sequencing and linkage analysis of SNPs located nearby the gene of interest provide a convenient and efficient way for PGD experiment design. For this couple, 116 informative SNPs (range: 83-292) were identified for each haplotype on average. The estimated ADO rate of each embryo was ranged from 0% to 31.71%. However, due to plenty of SNPs, there were still enough SNPs remained for haplotype analysis for embryos having a high ADO rate. Informative SNPs were distributed from upstream of *USH2A* gene to downstream of *USH2A* gene. This can guarantee the observation of recombination and reduce the error that may be caused by recombination if it occurs. The genotype was successfully determined for each embryo. In embryo 7, a recombination event was observed in the maternal allele, but it didn’t influence the genotype deduction of the *USH2A* gene because the recombination loci were outside the gene region. In this study, we simply captured 1.5 Mb region containing gene region and 350kb flanking region of *USH2A*. The capture region could be reduced by selecting highly heterozygous SNPs using public databases such as HapMap, dbSNP, 1000 Genome, et al., which can be helpful in reducing the cost of sequencing.

Invasive prenatal diagnosis is an important approach to make sure that the fetus is healthy during PGD cycle. On the one hand, analyzing mutations of interest directly withfetal genomic DNA can further confirm the accuracy of IVF-PGD procedure. On the other hand, it has been reported that the IVF embryo may have a higher risk of chromosomal anomaly than spontaneous pregnancy, and the embryo may be chimeric in some cases[12]. This means that the genetic make-up of biopsied cells may be different from the fetus, so invasive prenatal testing for chromosome anomaly is very important. Recently, it has been proved that low-coverage whole-genome NGS is a sensitive and high-resolution method to detect chromosomal aneuploidy and large imbalanced arrangements. It can detect 25% of mosaisic chromosmal anomally [7]. Thus, performing an NGS-based invasive prenatal chromosome abnormality detection can provide important genetic information.

## 5 Conclusions

In conclusion, we here present a procedure combining MPS-based PGD and invasive prenatal chromosomal anomaly detection in an Usher syndrome-risked family and have obtained a successful outcome. We believe that the capture MPS is a powerful tool for developing personalized PGD diagnosis which can be easily extended to other genes. And the integrated application of different MPS-based genetic detection methods in various reproductive stages can provide comprehensive information for genetic counseling and clinical decision.

## Compliance with Ethical Standards

### Funding

The study was carried out with the fund support of Tianjin Key Technologies Research and Development Program (Program Number: 05YFGZGX09900) and Major Technical Innovation Project of Hubei Province (Program Number: 2017ACA097).

### Informed consent

Informed consent was obtained from all patients included in the study.

### Ethical approval

All procedures performed in studies involving human participants were in accordance with the ethical standards of the institutional and/or national research committee and with the 1964 Helsinki declaration and its later amendments or comparable ethical standards.

---

**Supplemental Fig. 1.**
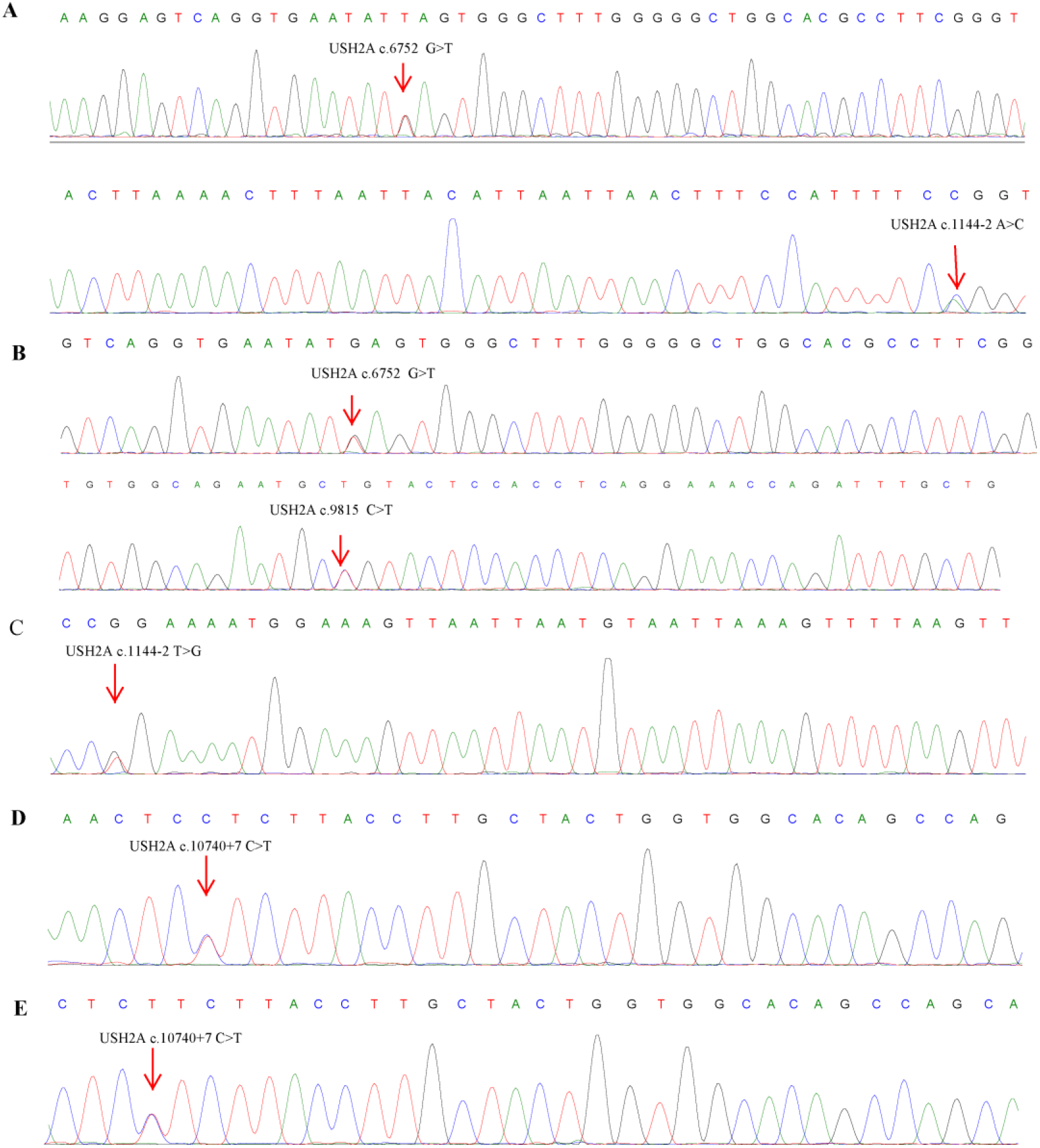
Sanger sequencing result of each family member. a. The result of the patient showed heterozygous of c.6752C>A mutation and c.1144-2A>C mutation in *USH2A*. b. The result of the patient’s father showed heterozygous of c.6752C>A mutation and 9815C>T mutation in *USH2A.* c. The result of the patient’s mother showed heterozygous of c.1144-2A>C mutation in *USH2A*. d. The result of the patient’s husband showed heterozygous of c.10740+7 G>A mutation in *USH2A*. e. The result of the patient’s father in law showed heterozygous of c.10740+7 G>A mutation in *USH2A*

**Supplemental Fig. 2.**
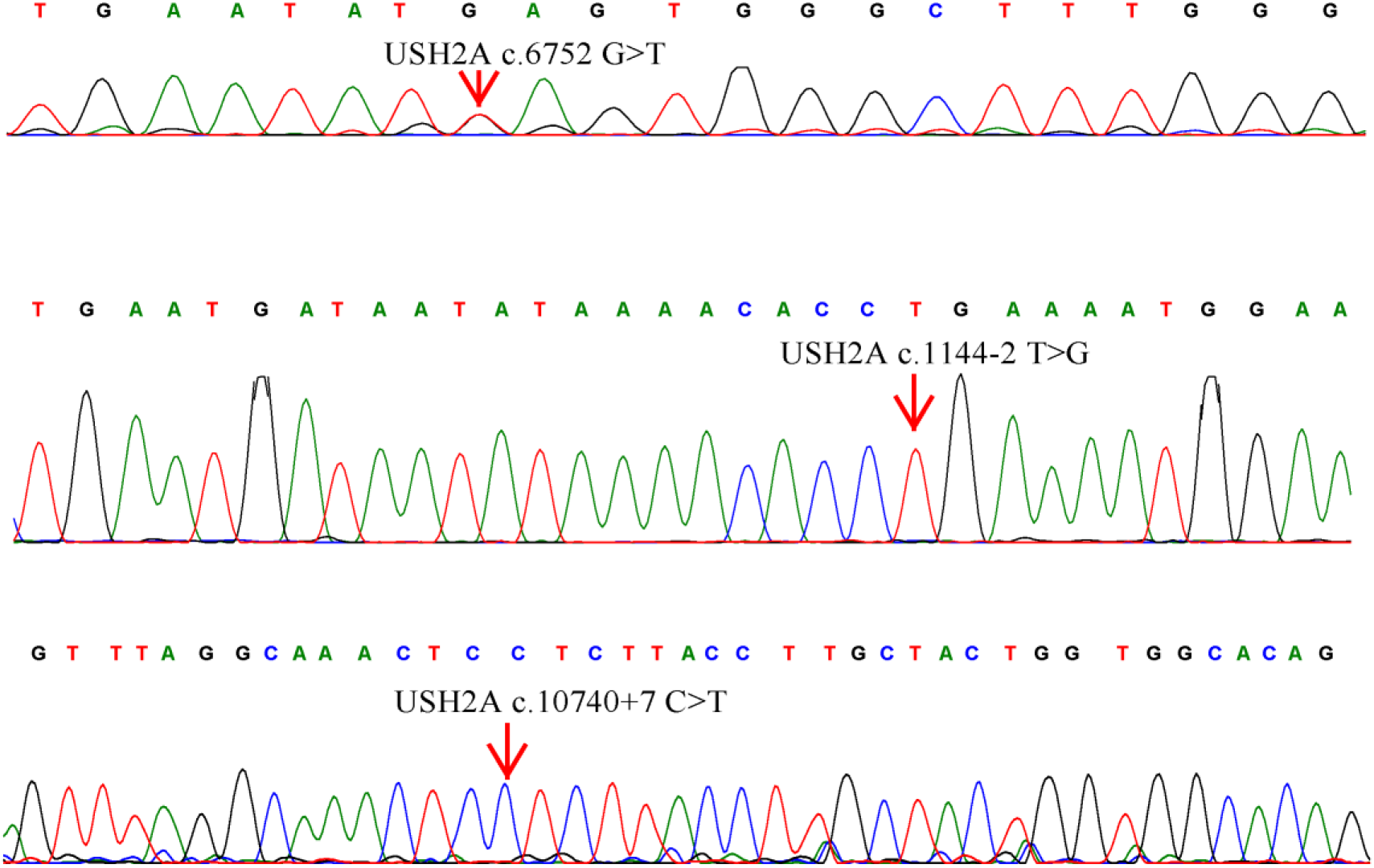
Sanger sequencing results of the baby’s cord blood. The results showed the heterozygous c.6752C>A mutation and wild type of c.1144-2A and c.10740+7G in *USH2A*

